# Pulling water from a stone: subaerial biofilm water dynamics on porous solid materials

**DOI:** 10.64898/2026.10.09.758030

**Authors:** Fabiana Russo, Isaac Klapper

**Affiliations:** Department of Mathematics and Applications “Renato Caccioppoli”, University of Naples Federico II; Department of Mathematics, Temple University, Philadelphia, PA, USA

**Keywords:** Subaerial biofilm, water activity, mathematical model, model validation

## Abstract

Subaerial biofilms (SABs) are microbial communities that colonize air-exposed surfaces in both natural and engineered environments, forming an interface between the atmosphere and the substrate. Often water availability is crucial for their metabolic activity and survival. While precipitation may contribute to SAB hydration, the dynamic processes of evaporation and condensation – closely linked to fluctuating temperature and humidity conditions – can be key determinants of water availability and they constitute the main focus of this study. We present a model connecting the water-related dynamics in an SAB, its surrounding ambient air and underlying stone/masonry. A central feature of the model is the increased water affinity of the SAB in comparison to the stone pore network. Based on thermodynamic principles, the model captures changes in liquid water content and availability by accounting for the evaporation and condensation at liquid-gas interfaces through gradients in water chemical potential. It accounts for the SAB’s capacity to delay water loss by drawing water from substrate material pores to compensate for atmospheric evaporation. The proposed model is compared to prior experimental observations of water transport in limestone samples with and without SAB. Additional numerical simulations under variable environmental conditions demonstrate how permeability of the substrate layer may influence SAB water activity, and how the presence or absence of an SAB might affect water content within stone/masonry.

## 1 Introduction

Abiotic stone and masonry substrates exposed to the atmosphere, including both natural environments such as rocks and soils as well as engineered structures such as infrastructure and stone heritage, are often colonized by thin layers of densely packed microbes embedded in a matrix of extracellular polymeric substances (EPS) [1, 11, 12, 43]. These aggregates, commonly referred to as subaerial biofilms (SABs), form self-sustaining microbial ecosystems that can develop wherever direct exposure to the atmosphere occurs, despite often inhospitable conditions [43]. Survival in such environments is restricted to organisms capable of tolerating multiple and highly fluctuating environmental stresses. As a result, these communities are predominantly composed of stress-resilient fungi, microalgae, cyanobacteria, and heterotrophic bacteria [1, 11, 51].

Sessile aggregation offers enhanced protection against multiple stressors – including desiccation, nutrient scarcity, and antimicrobial agents – and facilitates microbial interactions and exchange of genetic material [11, 43]. SAB colonizers secrete extracellular polymeric substances that surround developing microcolonies, spread onto the substrate, and facilitate both cellular cohesion and adhesion to the surface [11, 52]. The formation of EPS-embedded microcolonies is a characteristic feature of biofilm growth [8]. However, despite structural similarities in their matrices, subaerial and submerged biofilms exhibit marked differences arising from the diverse environmental condition they encounter. In contrast to submerged biofilms, SABs have a more heterogeneous, patchy organization and can experience dramatic cyclic fluctuations in temperature and humidity [1, 12]. An example SAB can be seen in Figure 1.

**Figure 1.**
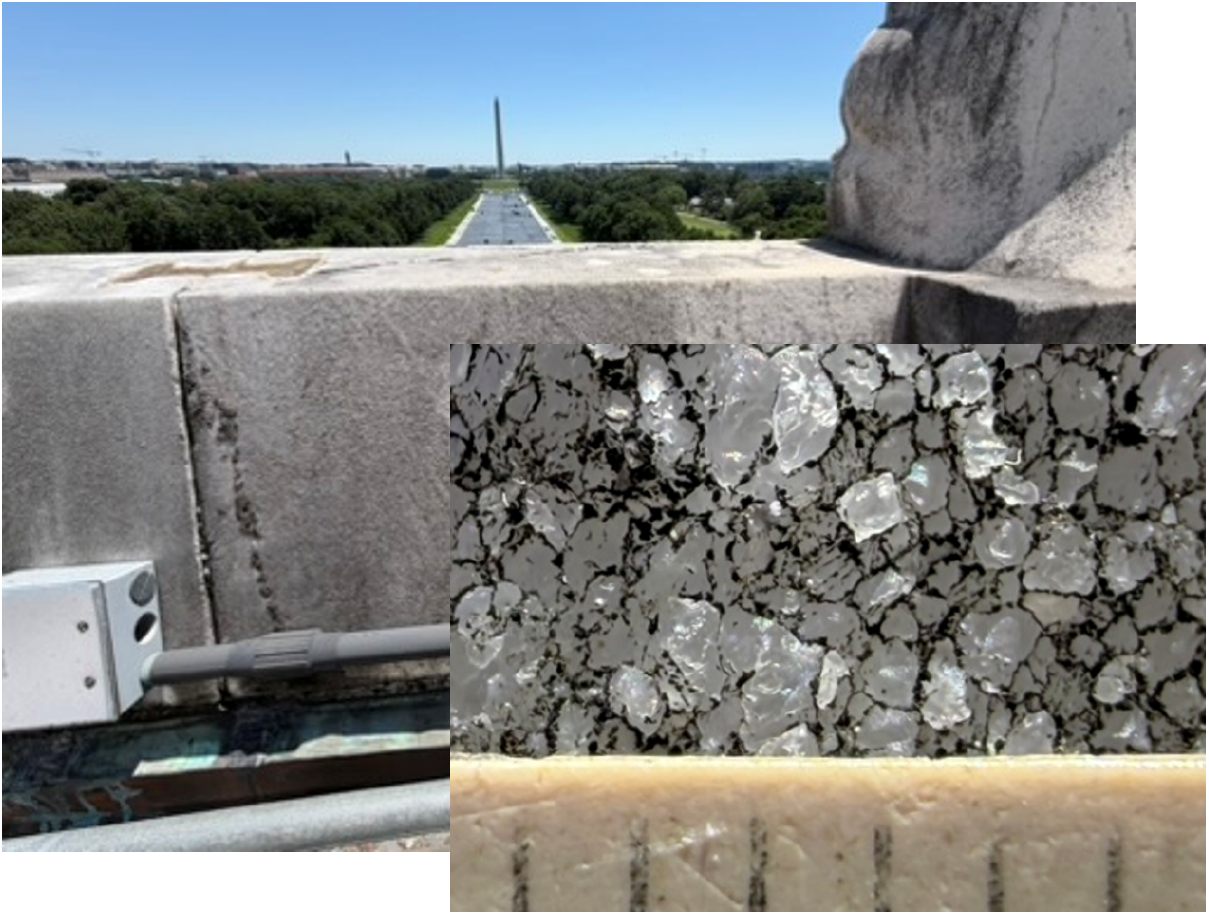
Macroscopic images of an SAB on marble, roof of the Lincoln Memorial, Washington D.C., at two observation scales: far and close-up views. Note SAB in between marble grains. Scale marks are 1mm apart. Photo credit: J. D. Grattepanche, Department of Biology, Temple University, Philadelphia, PA, USA.

SAB activity is regulated by multiple external drivers, including atmospheric humidity, solar radiation, and temperature, as well as the deposition of organic and inorganic matter [1, 11, 46, 51], with water availability emerging as a primary constraint on metabolic activity and survival in many environments [11, 21, 44]. Outdoor SABs typically rely on oxygenic phototrophs, such as cyanobacteria and algae, which convert inorganic carbon, i.e., CO_2_, into organic forms, supporting other microbes [1, 24, 41, 40, 46]. Notably, water, essential for cellular metabolic activities, structure, and function, is often in short supply in subaerial environments and, even when light and atmospheric CO_2_ are sufficient for photosynthesis, microbial activity is often particularly sensitive to water availability [11, 10, 13, 21, 23, 40, 49].

Water inputs may derive not only from precipitation, but also from dew, fog, and evaporation/condensation [1, 12, 21, 41]. Stone/masonry substrate surface and interior architecture and water retention capacity emerge as possibly important factors shaping the composition of subaerial microbial communities. The substrate not only provides mechanical stability to microbial communities, but, as a porous material, may modulate water availability by functioning as reservoir and buffer for moisture [1, 21, 49, 46, 43]. Natural rock and engineered porous materials are typically composed of a background phase, commonly referred to as matrix, interspersed with pores and voids that vary widely in size, shape, and spatial arrangement, and may be either isolated or interconnected. Porosity and pore structure are fundamental in governing the permeation, transport, and storage of water [3, 18]. They depend, for example, on mineral composition and formation history of the rock and can subsequently be altered by processes such as deformation and weathering [3, 10]. In particular, it has been posited that surface heterogeneities, such as pores, fissures, and cracks, generate favorable micro-environmental conditions in which microorganisms benefit from localized moisture accumulation [11, 21, 46]. Also, EPS matrix may play a role in water storage: beyond providing structural support and protection, EPS is believed sometimes to retain water, enabling prolonged metabolic activity and providing protection under desiccating conditions [1, 4, 11, 43, 44, 52]. Notably here, we propose that EPS may also act as an interface barrier, limiting and retarding water loss from the stone/masonry to the air [20, 34, 35, 44].

External conditions can not only affect SAB internal organization but also alter interactions with both the underlying solid substrate and the surrounding atmosphere. Environmental factors such as temperature and humidity may be fundamental in regulating water balance within the whole ecosystem – composed of the SAB and the substrate – influencing biofilm hydration and, consequently, microbial activity, mineral dissolution, and nutrient cycling. In particular, temperature and humidity conditions ultimately govern evaporation and condensation processes, which in turn control the overall exchange of moisture between an SAB, its underlying material, and the surrounding environment. Understanding these processes is therefore essential for capturing water dynamics in both natural and engineered environments, and for studying the role of water in these ecosystems, as it has the potential to influence energy production, nutrient exchange, and contribute to maintenence of SAB community life under desiccation. Analyzing physiological responses of SABs and substrate to water-related dynamics provides insight into their adaptive mechanisms and the evolving biofilm-stone relationship, which forms the motivation for this study.

Despite their relevance, the nature of the interactions linking an air-exposed substrate to its overlying SAB community remain understudied. Over the past decades, mathematical modeling has been employed to study biofilm development and microbial interactions, including communities composed of photosynthetic and heterotrophic microorganisms [27, 28, 32, 38, 39, 50]. However, the majority of existing models have been developed for submerged biofilms or biofilms on agar [47], where water availability is not limiting and/or interactions with the supporting substratum are usually neglected. This modeling framework is less suited to describe systems in which water limitations and substrate-biofilm interactions become significant. In response, a mathematical model has been proposed for predicting SAB water activity based on external temperature and relative humidity conditions [41] as well as a modeling framework to investigate the interplay between cyanobacteria and heterotrophic bacteria, accounting for their dependence on local environmental conditions such as temperature, humidity, light intensity, and environmental CO_2_ [37, 40]. However, these models neglect interactions between the SAB and substrate, which is treated as impermeable. Consequently, the influence of dynamic water exchange on SAB hydration levels and water availability remains largely unexplored. The present study thus aims to develop a mathematical description of water-related dynamics in a system composed of an SAB and the underlying solid substrate by identifying key liquid-vapor interfaces and incorporating relevant thermodynamic principles.

To do so, a system of equations is introduced to describe temporal variations in content of liquid water within both the stone/masonry and the SAB as a result of evaporation and condensation processes occurring within each layer, as well as the water exchange between them and the atmosphere. Stone/masonry is represented as a porous medium composed of interconnected spherical pores partially filled with water, while the SAB is modeled as a thin structure that may behave either as a non-porous or a porous medium, depending on its hydration state. A key feature of the model is the affinity of both substrate pore network and SAB matrix for water, which governs water availability and mobility in the two compartments. In particular, SABs are posited to have the greater affinity, in a sense defined below. In the following, water transport and accessibility within stone/masonry and SAB are formulated as functions of their respective water saturations. In addition, the model accounts for the ability of SABs, at high water content, to insulate interactions between the underlying substrate and the surrounding atmosphere, thereby effecting exchange of water. This mechanism leads to the identification of two distinct regimes for the substrate: *closed environment* and *open environment*. In the closed regime, air within the pores has no direct connection with the external environment, and evaporation and condensation processes are strongly constrained. In contrast, in the open regime, pore air remains connected to the outside atmosphere, facilitating moisture exchange and evaporation-condensation dynamics. In this way, an SAB, when hydrated, acts as a barrier that isolates the stone/masonry surface and delays water evaporation, while in a desiccated state, moisture transport from atmosphere to substrate is allowed.

Central components of the proposed model are supported by comparing predictions with experimental data reported in the literature. Particular attention is given to results presented in [44], where water evaporation was observed and measured from limestone samples either SAB-covered or SAB-free, under controlled temperature and humidity conditions. This comparison provides support for the model and allows the estimation of some key model parameters. Subsequently, numerical simulations are performed under variable environmental conditions featuring daily fluctuations to assess the importance of evaporation and condensation processes on water availability and mobility in porous stone/masonry ecosystems. Several scenarios are explored by considering different permeability conditions of the stone/masonry layer, and the presence or absence of an SAB.

## 2 Model definition

Evaporation and condensation processes, governed by nonequilibrium thermodynamics, involve the transport of water molecules across a liquid-vapor interface and are influenced by environmental factors such as temperature, pressure, and relative humidity. By identifying exchange mechanisms, a model will be described which captures water transport within a system composed of an SAB and a porous solid substrate like stone or masonry. (For brevity, and also because examples used are stone-related, we will refer to it as an SAB-stone system.) The model will account for variations in water content within both the SAB and the underlying solid layer and the interfacial exchange between them.

### 2.1 Evaporation and condensation

We rely on the Hertz-Knudsen equation, which is widely employed to describe evaporation and condensation processes at the interface between liquid and vapor phases [14, 15, 17, 19, 25]. When the substance in question is water and the equation is expressed in terms of intensive thermodynamic variables, it describes the relationship between the water mass flux *J*_*w*_ – defined as the mass of water evaporating/condensing per unit time per unit area – and pressure, in the form

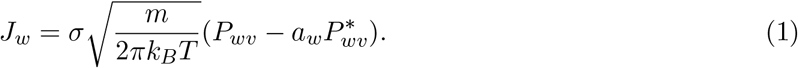

Here *m* is the mass of a water molecule, *k*_*B*_ is the Boltzmann constant, *T* is temperature, and *σ* is the evaporation or condensation coefficient, which quantifies the fraction of molecules that reach the interface and undergo a phase change. It is assumed that, near the interface, the temperatures of the liquid water and water vapor phases are equal, and that evaporation and condensation coefficients coincide. The term 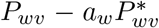 is the driving stress governing the evaporation and condensation processes, and its derivation is outlined in Appendix A. Pressure *P*_*wv*_ is the water vapor partial pressure, *a*_*w*_ is the water activity, which is discussed in the dedicated Section 2.3, and 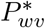 is the water vapor saturation pressure, which can be determined as described in Appendix B.

Net evaporation flux *J*_*w*_ is then the difference between the unidirectional evaporation flux and the unidirectional condensation flux [25]. When *J*_*w*_ *>* 0, condensation dominates, leading to an increase in the mass of liquid water in the liquid-vapor system. Conversely, when *J*_*w*_ *<* 0, evaporation prevails, resulting in a decrease in the system’s liquid water content. Mass of water evaporating/condensing in the system is related to changes in the liquid-vapor interface; the net evaporation rate *r*_*w*_ is given by

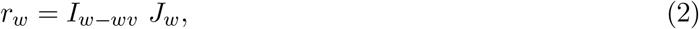

where *I*_*w−wv*_ depends on details of the interface between liquid water and water vapor undergoing the evaporation and condensation processes. See Section 2.2 for an application. The product *I*_*w−wv*_ *σ* sets the time scales for evaporation/condensation, which depend on material details, and are here regarded as empirically determined due to the complicated nature of the materials involved.

### 2.2 Evaporation and condensation in SAB-stone systems

In this section, the general Hertz-Knudsen equation (1) is adapted to describe water exchange mechanisms occurring in an SAB on a porous stone/masonry substrate, see Figure 2 for a rough schematic. We consider a three compartment model (ambient air, SAB, and porous material) and then study flow of water between compartments including dependencies on temperature and relative humidity. The stone/masonry layer is treated as a porous medium with interconnected pores which open to the overlying biofilm layer, and the biofilm layer is modeled as a thin continuous or porous medium, depending on its water content. Water exchange is assumed to occur at three interface types:

i. the interface between air and water within stone/masonry pores;
ii. the interface between air and SAB;
iii. the interface between SAB and stone/masonry pores.

**Figure 2.**
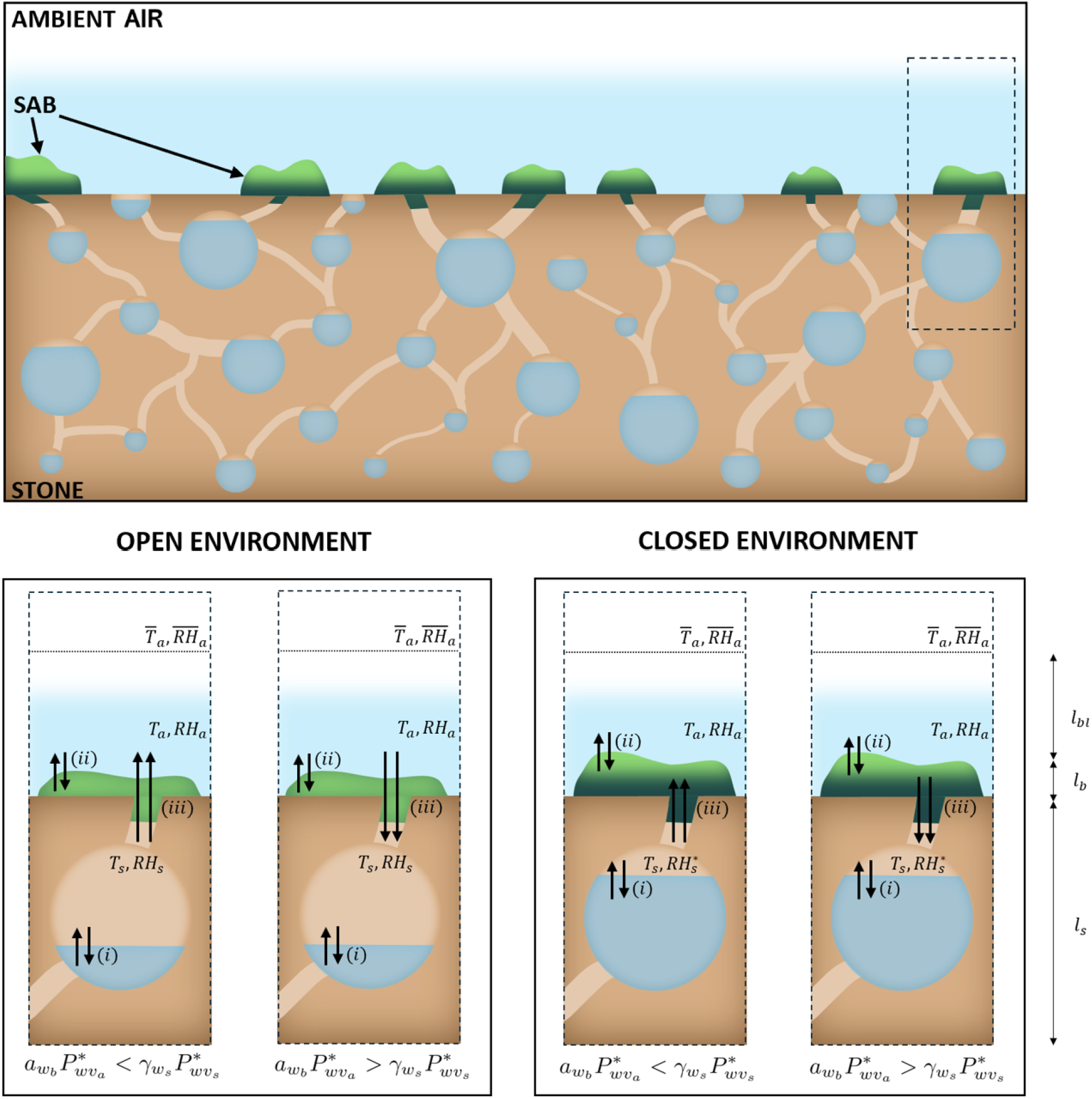
Schematic representation of the SAB-stone system. Depiction of the key exchange interfaces: between air and water in the stone/masonry pores (i); between external air and SAB (ii); between SAB and stone/masonry (iii). The SAB-stone system is shown under two different conditions: *open environment*, where the SAB is relatively poorly hydrated, allowing the stone’s superficial pores to remain connected to the surrounding environment; *closed environment*, where the SAB is relatively well hydrated, isolating those pores from the external ambient. Locations (i), *T*_*s*_ and *RH*_*s*_ denote temperature and relative humidity in the stone/masonry layer, which govern water evaporation/condensation at interface; locations (ii), *T*_*a*_ and *RH*_*a*_ indicate temperature and relative humidity in the parcel of air adjacent to the SAB, which regulate water exchange at interface; 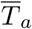 and 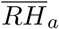 are temperature and relative humidity of the ambient air; locations (iii), all the environmental parameters influence water exchange between SAB and stone/masonry at interface. Relative humidity within the pores is denoted by *RH*_*s*_ when the system is in the open regime. In the closed regime, however, it differs from this value, and is denoted by 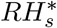 (water vapor partial pressure in the pores is evaluated using the ideal gas law).

Exchange fluxes at interfaces are primarily driven by variations in temperature and relative humidity according to equations of the form

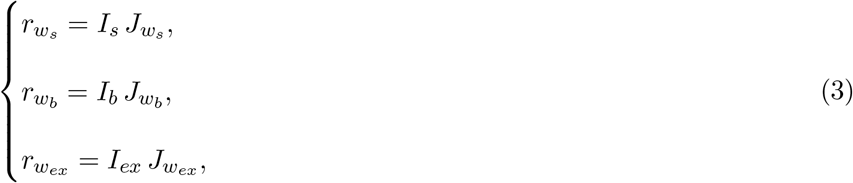

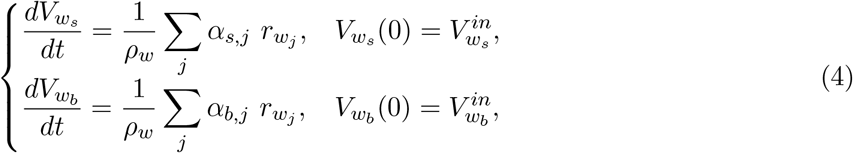

where subscripts *{s, b, ex}* refer respectively to the water exchanges between air and water in the stone/masonry pores, between air and SAB, or between SAB and stone/masonry, *J*_*w*_*j* and *r*_*w*_*j* denote the water exchange flux and rate occurring at interface *j* with *j* ∈ *{s, b, ex}*, and *I*_*j*_ is the area of the interface across which the exchange occurs. Water exchange rates *r*_*w*_*j* are governed by system (3), while variations in the volumes of liquid water in the stone/masonry *V*_*w*_*s* and SAB *V*_*w*_*b* are evaluated through system (4) where the right-hand sides account for the net rates of water evaporation, condensation or absorption occurring at the interface *j*, see schematic representation depicted in Figure 2. Water density is denoted by *ρ*_*w*_, while *α*_*s,j*_ and *α*_*b,j*_ are coefficients taking value from the set *{−*1, 0, 1*}* depending on whether the corresponding mass flux leads to a loss, no change, or a gain in water content for the respective layer. Specifically, *α*_*i,j*_ = 0 if layers *i* and *j* do not communicate directly.

#### 2.2.1 Water exchange within stone pores

We model rock/masonry as a matrix with interconnected, spherical-shaped pores, see Figure 2, borrowing a representation which has been commonly employed in both modeling and experimental studies, to describe for instance flow through porous media and to investigate the distribution of water within the pore space [9, 16, 48], as well as to evaluate material properties [3, 6]. Pores are assumed here to host two fluid phases: liquid water and water vapor. In the absence of a detailed knowledge of pore structure, the liquid-vapor interface within them is roughly approximated as 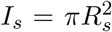, where *R*_*s*_ is the effective radius representing the combined volume of all spherical pores. This radius is given by *R*_*s*_ = (3*ϕ*_*s*_*V*_*s*_*/*4*π*)^1*/*3^, where *ϕ*_*s*_ is the stone porosity, defined as the ratio between the total pores volume *V*_*p*_*s* and the total volume of the porous medium *V*_*s*_ [36]. A more refined representation could be used with additional information about pore structure. Also, we note that, effectively, we are assuming that the liquid-vapor balance is uniform in the pore network, though in reality it would generally be spatially heterogeneous.

Temperature and relative humidity, which directly affect the dynamics of evaporation and condensation processes in the stone layer, are here denoted by *T*_*s*_ and *RH*_*s*_ (these are also assumed spatially homogeneous). To describe water dynamics within pores, Equation (2) becomes

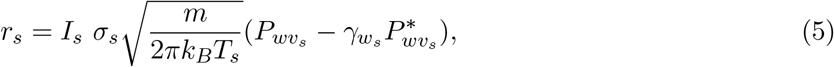

where 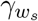 denotes the activity coefficient of water of the stone/masonry. We assume that liquid pore water can be considered either pure or a sufficiently diluted aqueous solution, so that the water mole fraction can be approximated as *N*_*w*_ = 1. Consequently, 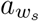 in Equations (1)-(2) coincides with 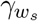; see Appendix A for further details. (Note that this is not always the case. As an example, salt contamination can occur.) The activity coefficient of water 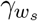 is modeled as a function of the water saturation in the pores, as detailed below in Section 2.3.

#### 2.2.2 Water flow and SABs

We suppose that the SAB layer exchanges water with both ambient air and underlying porous rock through an interface that encompasses its entire surface area *I*_*b*_. Similarly to the underlying porous media layer, recall Section 2.2.1, these exchange mechanisms are subject to environmental conditions in the parcel of air adjacent to the SAB: temperature *T*_*a*_ and relative humidity *RH*_*a*_. Influence of the thermal boundary layer is incorporated into the model to account for the temperature gradients resulting from heat exchange between the surface and the ambient air. Thickness of the thermal boundary layer depends on air thermal diffusivity and viscosity, as well as the flow conditions, including velocity and turbulence. This thin insulating layer between the stone surface and the ambient air modulates heat transfer, with increasing thickness leading to reduced heat exchange and larger temperature differences. The parameters *T*_*a*_ and *RH*_*a*_ at the interface biofilm-boundary layer can be computed from the environmental conditions in the stone and in the ambient air using the relationship proposed in [41] and recalled for convenience in Appendix B.

We propose here that water in the porous rock can serve as a water reservoir for the SAB [21, 46]. That is, water stored within the porous stone can migrate outward toward the SAB, where it may be absorbed and utilized. This transfer occurs, during periods of relatively low ambient air *RH*_*a*_, as SAB draw water from stone’s pores to compensate for atmospheric evaporation. As the biofilm absorbs this migrating water, higher water activity can be temporarily maintained, allowing microbial processes to continue despite evaporative loss of water to the atmosphere [44].

To describe water dynamics in the SAB, rate (2) becomes rates

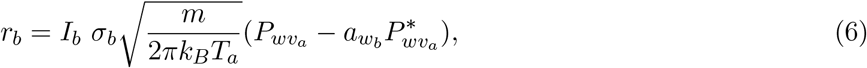

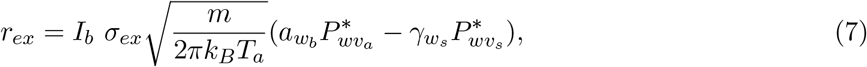

which describe water transport within the biofilm occurring at the interface between the SAB and the ambient air and the water exchange occurring at the interface between the stone/masonry and the SAB. SAB water activity 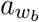 indicates, roughly, how readily available liquid water is for microbial growth and metabolic activity and depends on the SAB water saturation (as detailed in Section 2.3), and as such is a central output of this study. Importantly, under the assumption of impermeable substrate and assuming that evaporation and condensation effectively occur instantaneously, the SAB water activity 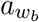 coincides with the relative humidity in the parcel of air adjacent to the SAB *RH*_*a*_, as proposed in [37, 40, 41].

### 2.3 Functional forms of the water activity and water activity coefficient

Water retention in solid materials depends on porosity, pore size, connectivity between pores, and specific material types [18]. In porous media composed of hydrophilic solids, such as clay, limestone, gypsum, and marble, water tends to adhere to solid surfaces and remain sorbed to the matrix [10, 33, 36]. Specifically, small amounts of water form thin films covering the particles of the solid matrix, and these strongly sorbed water molecules resist removal under normal conditions or, likewise, from being biologically utilized. The presence of subaerial biofilms, though, has been observed to be capable of influencing moisture dynamics and weathering processes. By modulating water exchange between the stone substrate and the surrounding environment, SABs may help maintain a more stable internal moisture content [34, 44, 46], thereby reducing hydraulic stress. Notably, it is hypothesized here that SABs – notably their EPS matrix [1, 11, 43, 52] – have the capacity to retain moisture not only from atmospheric humidity and precipitation, but also from the underlying substrate, acting as a sponge [1, 44, 46]. This retained moisture contributes to the biofilm’s hydration and overall functionality. On the other hand, water retained in such manners may not behave as an ideal solvent, particularly at lower liquid water volume fractions [18]. It is not immediately clear whether water pulled into an EPS network, for example, is readily available to nearby microbial cells.

The physical basis for all of this is thermodynamic, as described in equations (5) and (6). When water saturation is low, there are strong physical/chemical forces attracting water, whereas when water saturation is high, such forces are themselves saturated and no longer attractive [18, 30]. Water saturation itself is defined as the ratio between the volume of water within a medium and the total volume not occupied by the solid matrix [36]. We express it in terms of time-dependent variables as follows, where the temporal dependencies are made explicit for clarity,

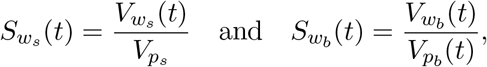

where 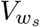 and 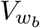 vary over time as a result of water exchange processes occurring within these media. The quantities 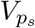 and 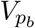 correspond, respectively, to the total volume of pores within the stone/masonry and the portion of the SAB volume that is not occupied by solid material.

Note that 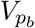 is a function of time, as observations suggest that the thickness of SABs varies according to their hydration state: hydrated SABs exhibit greater thickness than desiccated ones [42]. Indeed, it has been observed that certain laboratory-grown SABs, or those developing in highly humid environments, can reach thicknesses exceeding 50 *−* 100 *µm* [31, 45]. Consequently, stone pore volume 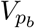 is here treated as a time-dependent variable, in contrast to 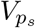, which is considered constant.

We set 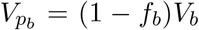, where *V*_*b*_ is the total volume of the SAB layer, and *f*_*b*_ denotes the biofilm solid fraction defined as the ratio between the constant volume of the solid matrix (consisting of solid components such as microbial cells, EPS, and others) and the total biofilm volume. The variability of the pore volumes within the SAB arises from changes in both the fraction of solid components *f*_*b*_ and the total biofilm volume *V*_*b*_; in particular, variations in the total volume are considered to result from changes in the SAB thickness *l*_*b*_. Here we assume that *l*_*b*_ varies from 50 *µm* (hydrated conditions) to 30 *µm* (desiccated conditions), while biofilm solid fraction *f*_*b*_ ranges from 50% to approximately 80% (value obtained considering that *l*_*b*_, and consequently *V*_*b*_, linearly varies with 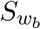) [42].

Water availability in the stone and SAB is quantified through water activity coefficient 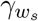 and water activity 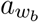, modeled as functions of the stone/masonry and SAB water saturation through the empirical constitutive relationships

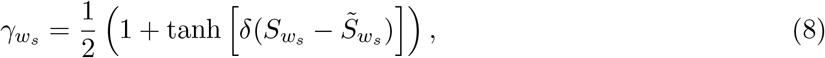

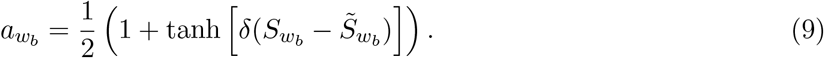

Here *δ* controls the sharpness of the transition, and 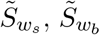 are threshold values indicating the onset of significant changes in 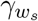 and 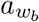, see Figure 3. These forms are chosen based on empirical observations of stone/masonry materials [18, 26, 36], with monotonic smooth transitions that vary between 0 and 1 as functions of 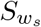 and 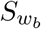, although details, e.g., transition points 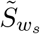 and 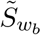, depend on the material properties. A material with a narrow grain-size distribution is often characterized by a sharp transition from saturated to unsaturated conditions, while the reduced number and size of pores in which the wetting fluid can reside, leads to a lower irreducible saturation (i.e., less water with high resistance to evaporation under normal climatic conditions) [26].

**Figure 3.**
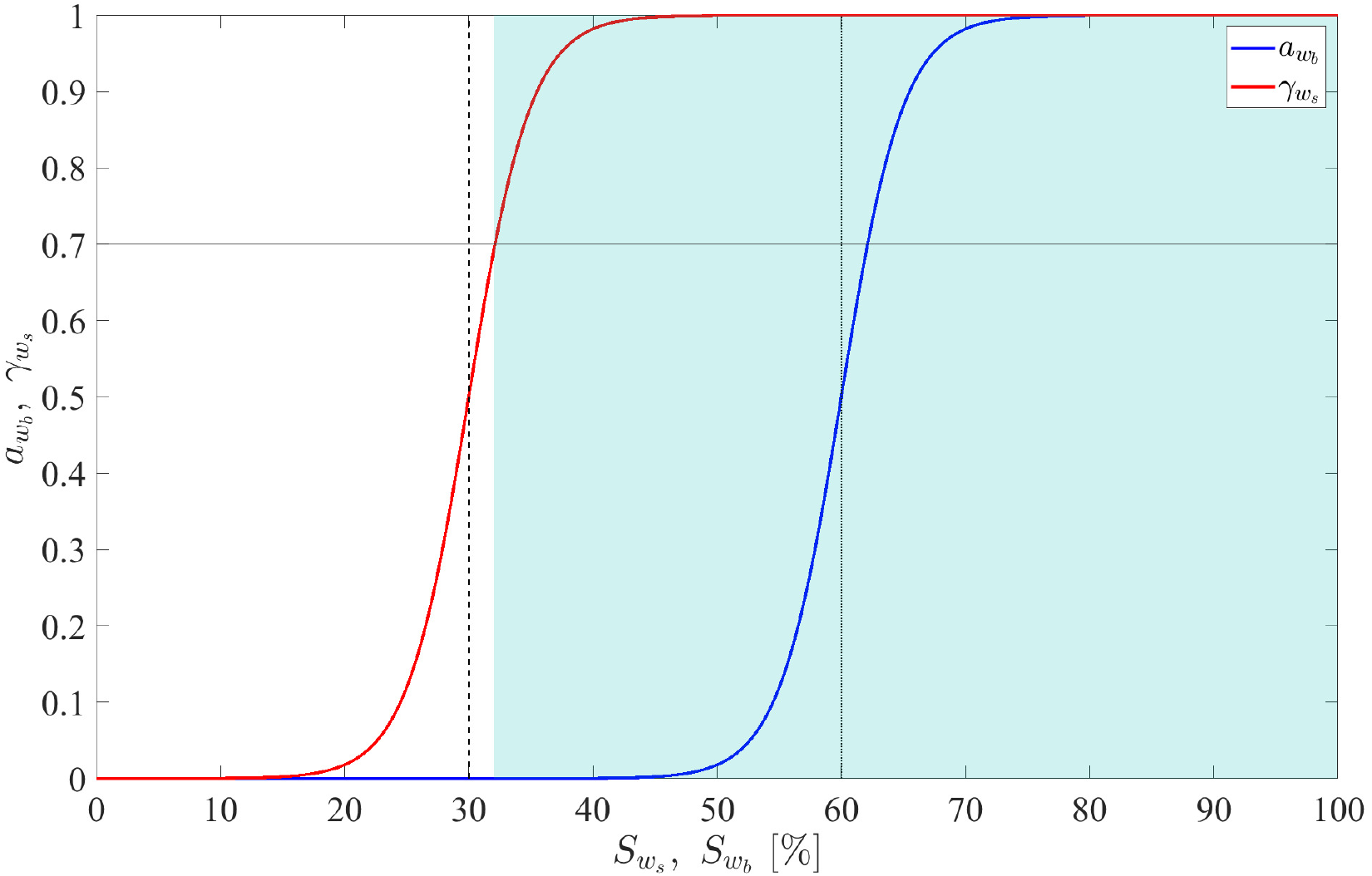
Stone pore water activity coefficient 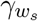 with respect to stone water saturation 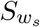(red) and SAB water activity 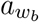 with respect SAB water saturation 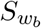 (blue), see Equations (8)-(9). Dashed and dotted black vertical lines individuate the water saturation thresholds for stone/masonry 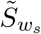 and SAB 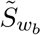, respectively. Referring to Equation (7) and noting that 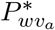 and 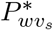 are approximately equal, then the rate of water transfer from stone/masonry to SAB is approximately proportional to 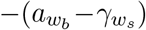. Experimental evidence from a lab SAB system [7, 44, 45] indicates that when the relative humidity exceeds approximately 70% (horizontal solid black line), the SAB rehydrated significantly and rapidly recovered metabolic activity. Thus, while stone/masonry water saturation and SAB water saturation are not in general equal, as long as stone water saturation is sufficiently large so that 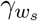 is close to 1 (i.e., the curve 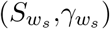 is in the light blue region), then the SAB can maintain relatively high water activity. Note that this observation is largely independent of the curve 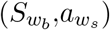, as long as its transition lies clearly to the right.

Larger values of *δ* produce a sharper transition, while smaller values yield a more gradual one. Here *δ* is set equal to 20. A relatively high value for *δ* is chosen because changes in activity have been observed to occur over a limited range of water saturations in porous media [18, 26]. This is also consistent with the observations on stone ecosystems reported in [44] (see Section 3 for a detailed analysis). Larger values of 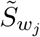 indicates stronger water affinity, i.e., “stickier” (to water) materials. Roughly, 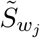 acts as a threshold separating surface-sorbed water from water that can participate in microbial activity. Note that for large *δ*: if 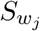 is lower than 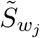, the quantities 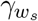 and 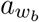 remain close to 0, indicating potential for a strong attractive driving force; conversely, when 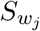 exceeds 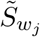, they are close to 1, corresponding to conditions where water is less strongly sorbed.

This discussion suggests a mechanism by which a wet SAB may maintain high water activity during drying atmospheric conditions. As an SAB dries, and its water saturation drops below 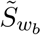, it can pull water from the underlying porous material which still has saturation above 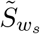, see Figure 3. The threshold value 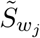 is set equal to 30% for the stone/masonry layer and 60% for the SAB layer, see Section 3 for justification. A high threshold value 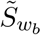 for SABs is not at first glance advantageous; it seemingly would necessitate higher relative humidity before SAB water activity can increase. However, we argue that, in combination with lower 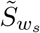, transiently large SAB water activity is possible. Note that this mechanism is relatively insensitive to the value of 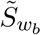 as long as it is sufficiently larger than 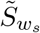.

### 2.4 Closed and open SAB conditions

A recent study [44] provides evidence of a dual-species laboratory SAB acting as a moisture-controlling layer, influencing water fluxes between its limestone substrate and environment. This positions the SAB as a mediator between the atmosphere and the stone, regulating to an extent water movement across the two layers [20, 34, 35, 44]. In light of this, and as an explanation, the proposed model, as illustrated in Figure 2, distinguishes between two distinct regimes for the stone pore network: a *closed environment* and an *open environment*. It is proposed here that when an SAB contains sufficient water, it effectively seals stone superficial pores, thus disconnecting its pore network from the surrounding air. In this closed system, the vapor phase within the pores quickly reaches saturation, i.e., relative humidity becomes 100%. Conversely, when an SAB desiccates, then its pores can communicate directly with the external air, enabling water exchange via vapor, i.e., water vapor partial pressure in the pores is approximately the same as that in the external ambient air (so that water content is also the same).

A transition between closed and open environment conditions is supposed to occur depending on a threshold value of the water activity 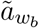. We set the value of this threshold to be somewhat higher than the water activity at which SABs are reported to become metabolically active, supposing that at this point there is enough liquid water to hydrate the entire SAB [7, 44, 45], and so isolate the underlying stone from the external environment. In particular, a value of 0.8 is assigned to this parameter. When the SAB water activity is below 0.8, the SAB is relatively dry, allowing moisture from the air to reach the porous medium. In this condition, water exchange between the stone and the atmosphere occurs freely, independently of an overlying SAB layer. However, when SAB water activity exceeds 0.8, the SAB is considered sufficiently hydrated, and further direct communication between stone pores and air is no longer allowed in the model. This marks the transition to a closed environment condition, where the stone becomes isolated in terms of moisture exchange.

We do not in fact apply a discontinuous transition. Rather, a function *η* is introduced to describe a smooth change from closed to open conditions, and is defined as a function of the SAB water activity by

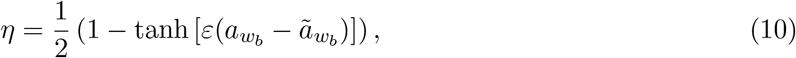

with *ε* = 15 employed in computations. The parameter *η* is defined using the same empirical functional form previously adopted for the water activity coefficient 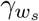 and the water activity 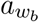, with a change in sign due to the opposite physical trend. For desiccated SABs, i.e., for low values of the water activity 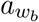, the value of *η* is close to 1. As hydration increases, *η* decreases, trending towards *closed environment* conditions. For highly hydrated SABs, i.e., for high values of 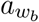, note that *η* effectively vanishes. In the absence of SABs, *η* coincides with 1, meaning that no biofilm-induced effect is present in the evaporation/condensation process within the stone pore network.

In light of the biological framework described, the model equations and appropriate initial conditions for modeling the dynamics of water in the SAB-stone system, as well as the liquid water content of both layers are

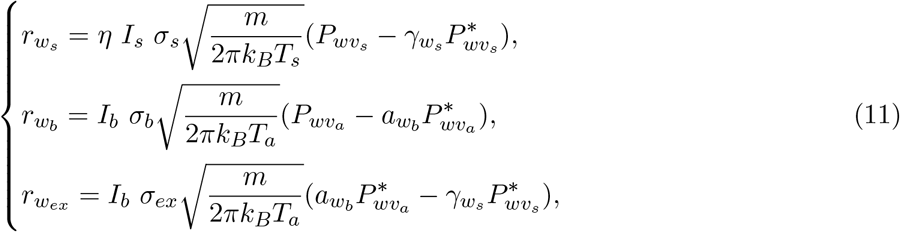

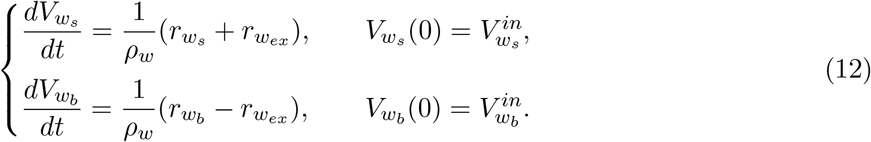

Pore water vapor partial pressure in the stone 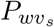 is given by

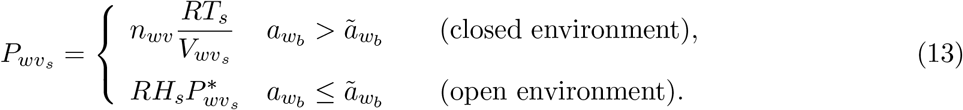

In the closed environment, the pore network behaves like a closed system and the water vapor partial pressure 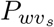 is determined using the ideal gas law. Once relative humidity reaches 100%, the vapor phase within the pores becomes saturated, effectively halting evaporation (or condensation). In open environment conditions, there is a direct connection between the network and the surrounding environment because pores are no longer sealed.

Equation (12)_1_ tracks variation of water volume in the stone pores, with rate determined by both the water exchange with the pore vapor phase and water in the SAB. Equation (12)_2_ describes water volume dynamics in the SAB, which depend on water exchange both with the external environment and with water in the stone. The water absorption term represents water transfer from SAB to stone, and so appears with a positive sign in the first equation and negative sign in the second.

System (11) shows that the interplay between temperature and humidity can either amplify or deamplify water exchange processes: during evaporation, higher temperatures or lower relative humidity enhance liquid to vapor transfer, while during condensation, lower temperatures or higher relative humidity promote water uptake into the stone pores or SAB. In addition to environmental factors, it is worth noting that the water content of both stone pores and SAB feeds back into shaping water dynamics. Indeed, activity coefficient *γ*_*w*_*s* and water activity *a*_*w*_*b* respectively depend directly on the saturation levels of the stone pores and the SAB (recall Section 2.3), thereby modulating both the intensity and direction of water transfer within the system as functions of current saturation.

## 3 Comparison to laboratory studies

Direct, *in vivo* observation of SABs is difficult at best, and laboratory measurements are also scarce; both are technically challenging. We refer here to [44], in which a series of experiments were conducted on dual-species autotrophic-heterotrophic SABs cultivated on solid stone substrates, and apply results to parameterize and test aspects of the mathematical model presented here. Particular attention is directed to observations of evaporation:

– water evaporation from an SAB-free limestone sample, used to determine *σ*_*s*_ in Equation (11)_1_, together with coefficients appearing in the functional form for 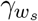, Equation (8).
– water evaporation from a limestone sample with SAB, used to determine *σ*_*b*_ and *σ*_*ex*_ in Equations (11)_2_ and (11)_3_, together with coefficients appearing in the functional forms for 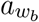 and *η*, Equations (9) and (10).

In [44], samples, after being saturated by immersion in liquid water, were placed abruptly in a climatic chamber, set at 25°*C* and 50% relative humidity, until moisture equilibrium with the environment was reached. The authors monitored over time the evaporation process of limestone specimens with and without desiccated SAB using thermography, which measures the cooling effect of latent energy release during the liquid-vapor phase transition, allowing evaporation flux to be inferred. These observations were further corroborated by direct evaporation flux measurements. Some of the experimental results presented and discussed in [44] are reprinted in Figure 4a and Figure 5a.

**Figure 4.**
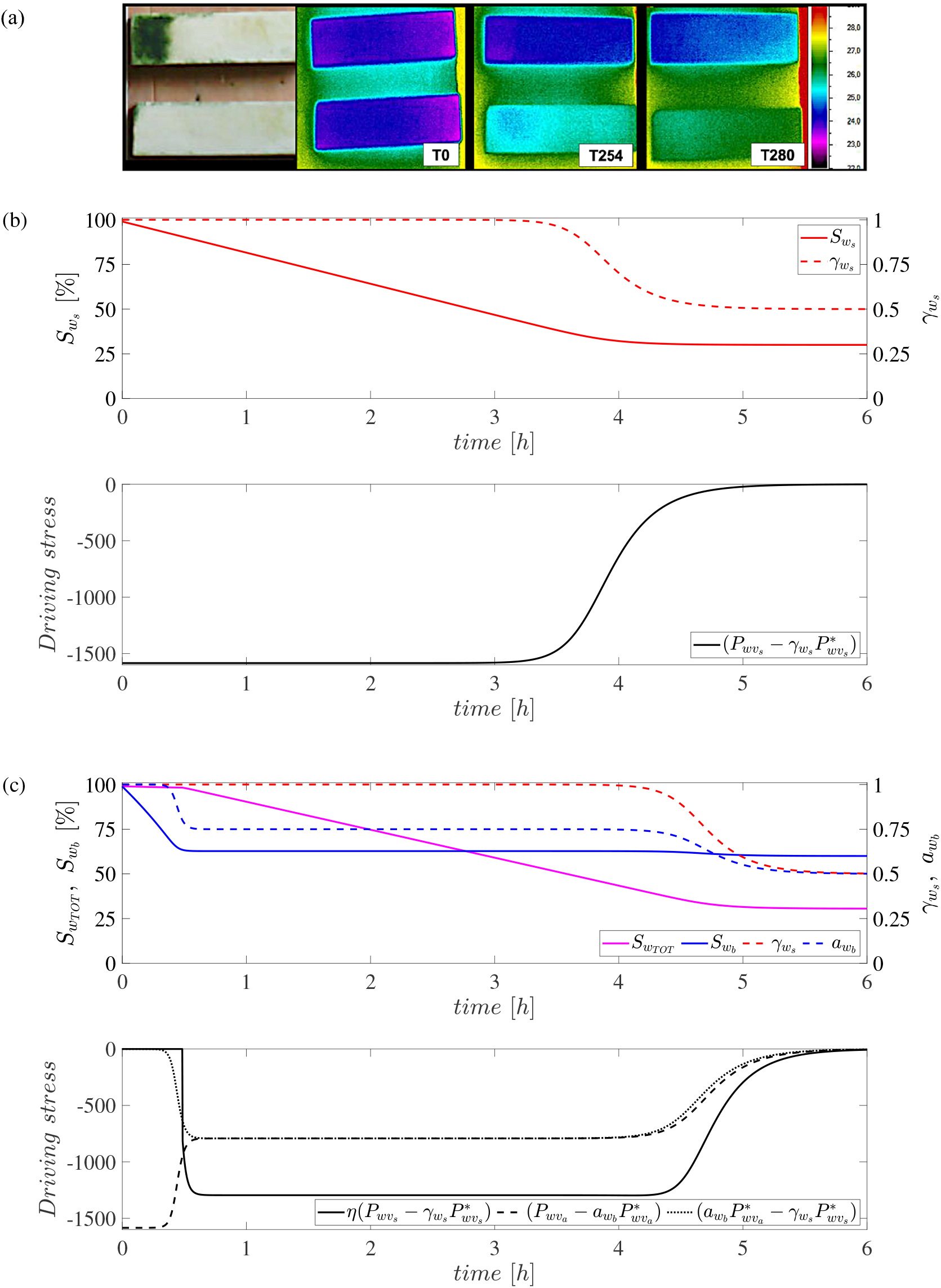
(a) From [44], reproduced under CC BY license. Commercial limestone specimens, composed of approximately 90% calcite and dolomite, were inoculated with a planktonic phototroph-heterotroph culture, resulting in development of a phototrophic SAB, and then placed in a climatic chamber. Water content was tracked until moisture equilibrium with the environment was reached. Thermography indicates evaporation (blue is cooler) for limestone samples with (top) and without (bottom) SAB at 0, 254, and 280 minutes. (b) Model results for comparison (without SAB): stone pore water activity coefficient 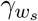, stone water saturation 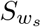, and the evaporation driving term in the SAB-free sample. Evaporation proceeds until water saturation drops below the threshold 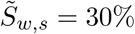 (at approximately 3.5 h), at which point the driving stress reduces towards 0 and evaporation slows. (c) Model results for comparison (with SAB): stone pore water activity coefficient 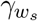, SAB water activity 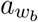, total water saturation 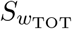, SAB water saturation 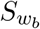 and the exchange driving terms in the SAB-covered sample. Note delay in evaporation drop (to approximately 4.5 h). Between approximately *t* = 0.5 h and *t* = 4.5 h, the SAB opens up and evaporation from the SAB to the ambient air is compensated by capture of water vapor from the underlying substrate.

**Figure 5.**
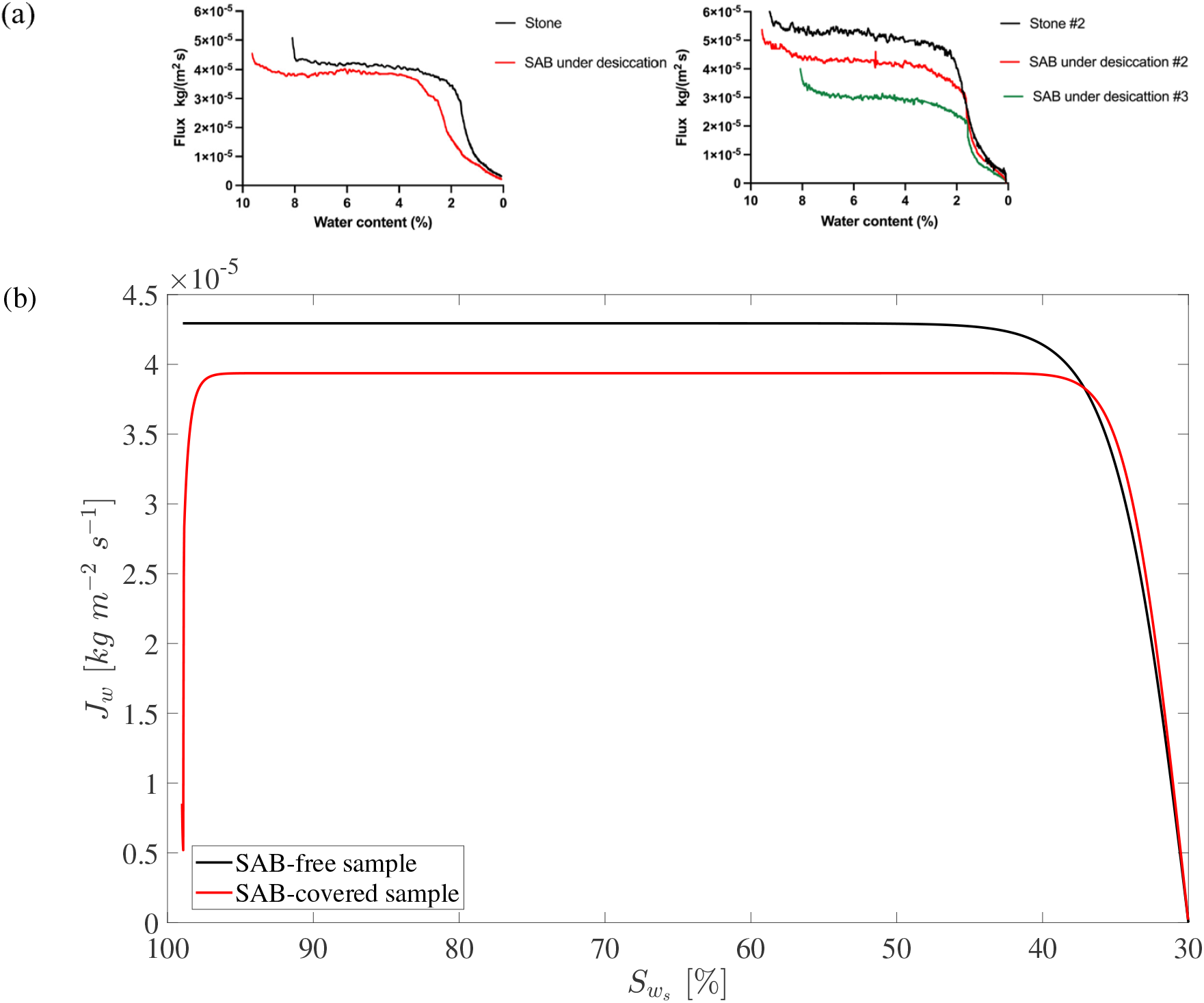
(a) Results from [44] and supplementary material, reproduced under CC BY license, using commercial limestone specimens composed of approximately 90% calcite and dolomite, with a porosity of approximately 8.91%. Samples were innoculated with a planktonic phototroph-heterotroph culture, resulting in development of a phototrophic SAB, and then placed in a climatic chamber until moisture equilibrium with the environment was reached. Measured evaporative flux of samples with and without SAB versus water content are shown. (b) Model simulation of this experiment: variation of computed absolute value of evaporation flux *J*_*w*_ with respect stone water saturation 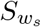 in the sample with and without SAB. Initial transient in the SAB-covered sample curve is due to initial conditions.

In the following, we simulate these experiments with two main goals. First, we look to set time scale parameters based on time scales observed in the experiments, and to set *η* via flux amplitudes. Note that coefficients *σ*_*j*_ essentially scale time, that is, they determine the time scales for water exchange processes. These are likely material-sensitive and are empirically determined here, as timescale information is relatively easily available. The other important class of data we have available from [44] are water fluxes. In particular, the difference in flux between SAB-covered and SAB-free cases allows estimation of transition function *η*, which becomes active during evaporation events and in presence of SAB coverage. Other parameters are either fixed by prior knowledge 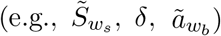 or hypothesized 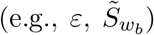. In the latter case, these parameters are related to “stickiness” properties, which are less easily measured directly. The central hypothesis, though, is that SABs need to be “stickier” than their underlying substrate, and we are able to test this qualitatively against measurements from [44], see below. Physical results, it turns out, are relatively insensitive to the degree of stickiness.

Once parameter choices have been made, as a second goal of the experimental comparisons, we wish to test key model suppositions: can the model reproduce central qualitative experimental results? The aim is to demonstrate that a relatively “sticky” biofilm layer can offer an explanation for water retention phenomena observed experimentally.

Model equations are integrated using an in-house MATLAB code based on the solver ‘ode15s’ and the simulations are carried out with the same environmental conditions, stone specimen dimensions, and porosity characteristics as those used in the experimental study (stone samples of dimension 75 *mm* x 23 *mm* x 10 *mm* were used, at set at 25^*°*^*C* and 50% relative humidity). Note: since relevant environmental details of the system were not reported in the experimental results, the effect of the thermal boundary layer (in the air above the SAB) is neglected when comparing numerical and experimental results.

### 3.1 Water evaporation in SAB-free and SAB-covered samples

To reproduce the experimental data in the case of SAB-free samples, model simulations were performed by considering equations (11)_1_ and (12)_1_ with *η* = 1 and *σ*_*ex*_ = 0 (i.e., effectively, absence of SAB). In conjunction, to simulate the SAB-covered case, the full system of Equations (11) and (12), with *η* as defined in (10), were considered. Equations in system (12) were assigned fully saturated initial conditions (i.e., water volume in the stone 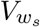 and in the SAB 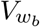 equal to the pore volume in the corresponding layer, respectively 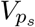 and 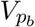). Results for SAB-free sample simulations are reported in Figure 4b and Figure 5b, while those for SAB-covered samples are shown in Figures 4c and Figure 5b. In the (model) SAB-free set-up, liquid water is free to evaporate directly from stone pores into the atmosphere, which proceeds over early times at an approximately linear rate, as long as water activity is high, see Figure 4b. Water activity remains high, recall Figure 3 for the relationship between water activity coefficient and water saturation, until sufficient evaporation has occurred so that saturation 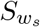drops to, approximately, threshold 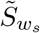. At lower saturations, pore stickiness sets in, reducing evaporation rate until, in fact, an equilibrium is approached. The coefficient *σ*_*s*_, which effectively determines the time scale, is set to 2.7 *·* 10^*−*4^, to match the approximately five-hour period to reach equilibrium observed for SAB-free samples in [44].

Figure 5b (black line) shows the computed absolute value of flux *J*_*w*_ with respect the stone water saturation, which is in agreement with the observed evaporative flux measurements reported in Figure 5a (black lines), providing a measure of validation, particularly of the form of the hypothesized pore water activity coefficient, recall (8) and Figure 3. It is important to note that [44] records the evaporation flux as a function of the water content of the sample, defined as the water mass contained in the sample with respect to the weight of the dried sample (oven 104^*°*^*C* for 24 h). In contrast, here the evaporation flux is plotted against the stone water saturation. Since the dry weights of the limestone samples used in the experiments are not available in [44], a direct comparison between these two curves is therefore not strictly possible (though it should be noted that the observed drop in flux, see Figure 5a, occurs at roughly 30% of maximum water content, matching the assumed form for 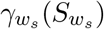, recall Figure 3, which can be seen as a measure of validation). Nevertheless, the computed flux curve is consistent with the experimental observations: the presence of a broad plateau indicates an approximately constant evaporative flux at higher water saturation levels, followed by a sharp decrease as the system approaches low saturation conditions. The abrupt change in flux is consistent with the functional form of the stone pore water activity coefficient, recall (8) and Figure 3. In both model and experiment, the maximum value of evaporation flux has an order of magnitude of 10^*−*5^ *kg m*^*−*2^ *s*^*−*1^.

The contrasting case of stone with SAB coverage is shown in Figure 4c. At early times, a transient evaporation loss to the ambient air occurs quickly from the very thin SAB layer, which, though buffered by the underlying stone, results in a rapid drop in its water content and, consequently, its water activity. There follows a long, pseudo-equilibrium state in which water transport from SAB to atmosphere is balanced by water transport from stone to SAB. During this period, the SAB is able to maintain an elevated water activity because, and only because, of water partially trapped in the stone. Once the excess water in the stone is depleted, at approximately *t* = 4.5 h, water activity drops to equilibrium with atmospheric conditions. Note also, in comparison to the SAB-free case, see Figure 4b, stone retains water longer, due to SAB evaporative insulation. Values of the *σ*_*b*_ and *σ*_*ex*_, which set time scales, are set to 5*·*10^*−*6^ to match the observed temporal evolution of water exchange in the SAB-stone system. Meanwhile, the threshold value 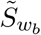 in relation (9) is set higher than the corresponding value for the bare stone, namely 60%. This choice is consistent with the different physical characteristics of the stone and the SABs and accounts for the higher biofilm stickiness, i.e., its stronger affinity for water. Again, we claim that results support the hypothesized form of the water activity, recall (9) and Figure 3.

Figure 6 shows the water mass contained in the model system for both SAB-free (solid line) and SAB-covered (dashed line) samples. Note that water content in the SAB-covered case decreases at a slower rate than in the SAB-free case due to evaporative insulation by the SAB. After approximately 4 hours, the SAB-free sample has already approached its minimum water content, whereas the SAB-covered sample still retains extra water for another hour, approximately. This behavior is consistent with the thermography images reported in Figure 4a.

**Figure 6.**
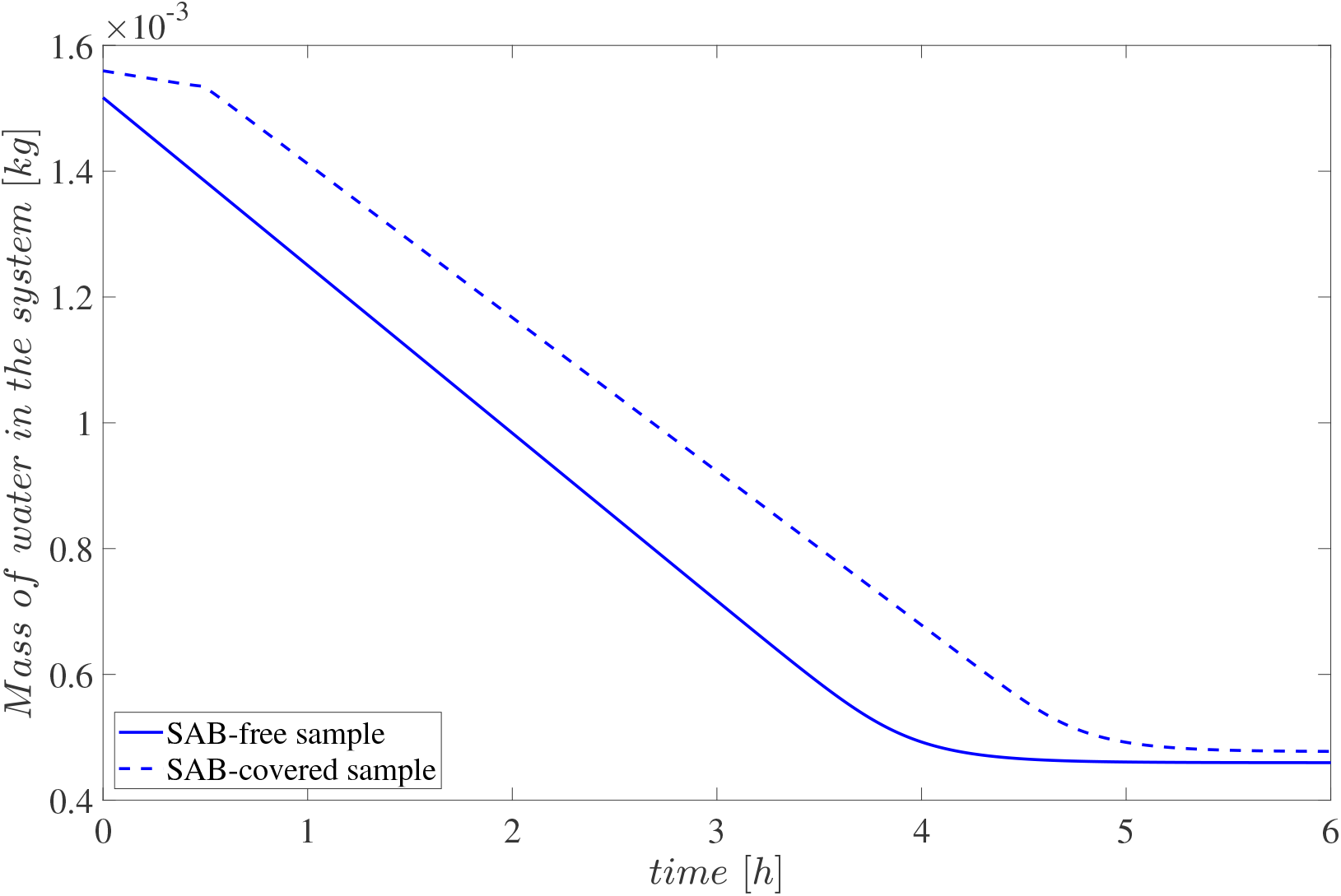
Total water mass contained in the (model) sample with and without SAB. SAB presence slows evaporation. The kink in the SAB-covered sample curve at approximately *t* = 0.5 hrs occurs when the SAB dries sufficiently so that stone pores become open to the ambient air.

### 3.2 Impact of stone/masonry on SABs

To emphasize the possible importance of water exchange from substrate to SAB we can constrain, in the model, the stone/masonry substrate to be effectively impermeable (so that it is unavailable as a water source for the SAB) by setting *σ*_*ex*_ = 0. Results are reported in Figure 7b where they can be compared with results in Figure 7a. Observe that, in the absence of water exchange with the stone, SAB water activity relatively quickly reaches equilibrium with the atmospheric RH. This behavior is due to the lack of water influx into the SAB, which would otherwise compensate for the evaporative loss (which is fast due to SAB thinness).

**Figure 7.**
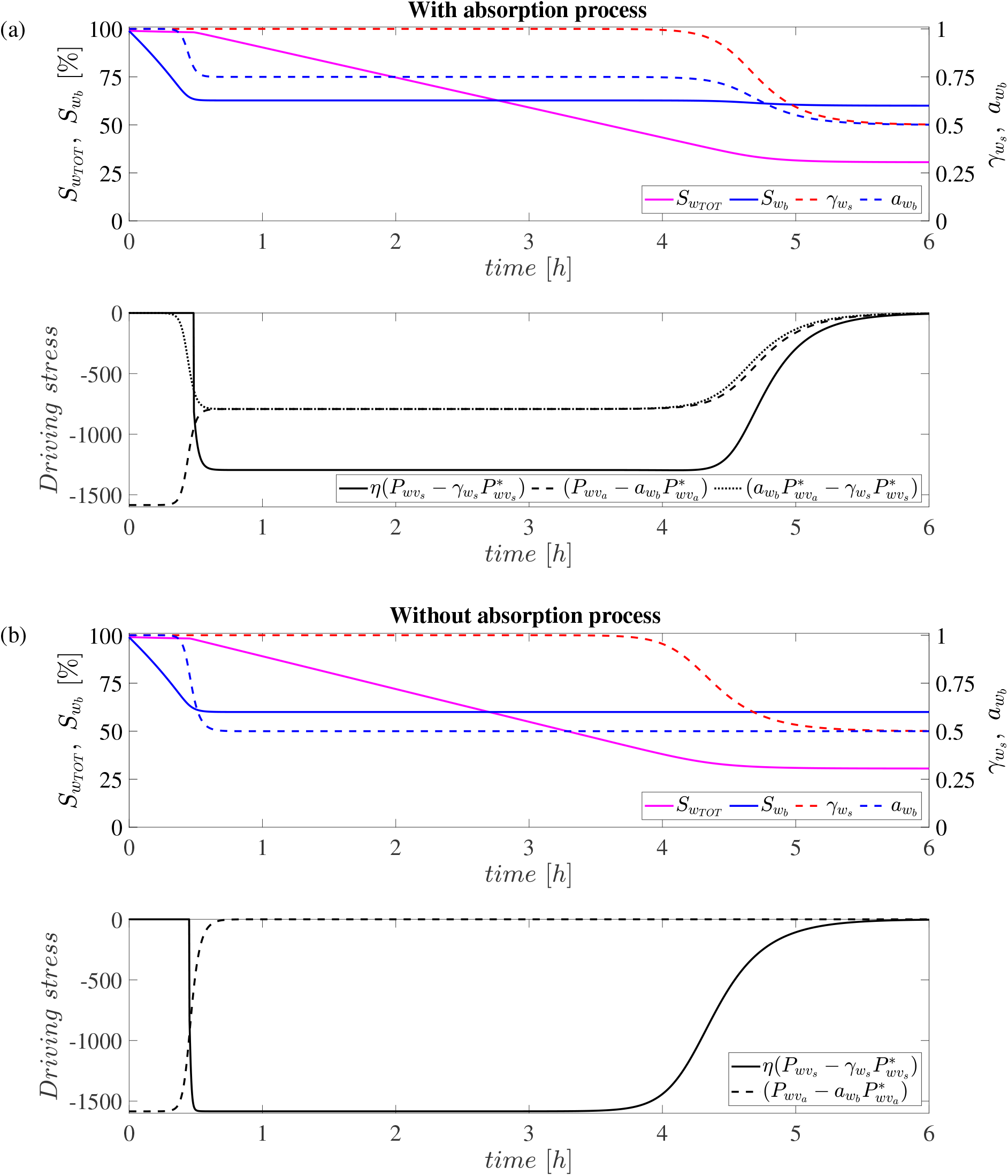
(a) SAB-covered sample accounting for SAB absorption: stone pore water activity coefficient 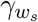, SAB water activity 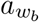, total water saturation 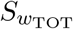, SAB water saturation 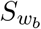 and the exchange driving terms. (b) SAB-covered sample neglecting SAB absorption: stone pore water activity coefficient 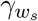, SAB water activity 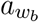, total water saturation 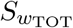, SAB water saturation 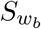 and the exchange driving terms. Note the difference in SAB water activity (blue dashed lines).

Note that even relatively small differences in SAB water activity can have significant biological consequences. The difference between water activity values of 0.5 and 0.75, which is not small, is the difference between viable and inviable conditions, but even changes in water activity of a few hundredths can be important metabolically [5]. A consequence of this observation is support for the hypothesis that stone/masonry water-related material properties may have important impact on SAB activity through an evaporation/condensation mechanism as described here.

## 4 Model study: water dynamics in SAB-stone systems under fluctuating environmental conditions

In Section 3, we compared model results to laboratory observations in order to calibrate evaporation/condensation time scale parameters and to provide a level of model verification. Those observations were conducted under controlled but artificial environmental conditions. To illustrate the possible role of SAB-stone water dynamics in environmental systems, simulations were performed using weather data collected on the marble roof of the portico at the Thomas Jefferson Memorial, Washington D.C. [41]. This dataset includes surface temperature *T*_*s*_ and relative humidity *RH*_*s*_ (measured directly on the substrate surface), ambient air temperature 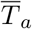(measured away from the stone surface), and light intensity. Ambient relative humidity 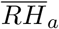 can be estimated following the approach described in [41], as recapitulated in Appendix B. The environmental parameter *T*_*a*_ and *RH*_*a*_, denoting the temperature and relative humidity in the parcel of air immediately adjacent to the biofilm, can be computed as also described in Appendix B. Since environmental conditions are input variables for the model, an average SAB thickness 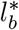 is used in the computation of *T*_*a*_ and *RH*_*a*_.

Simulations are based on the environmental profile of a typical summer day (June 18 *−* 19, 2016) at the Thomas Jefferson Memorial. Data for the 24-hour period from 06:00 AM, June 18, 2016, to 06:00 AM, June 19, 2016 are shown in Figure 8a. Background shading is correlated with light intensity. Note the day-night trend: temperatures drop and relative humidities rise at night, and vice-versa during daylight hours. Relative to the surrounding air, stone experiences more pronounced variations, which can be an important factor in SAB performance [11, 40]. Also note the abrupt drop in stone relative humidity occurring in late morning, consistent with the hypothesis of opening of SAB pores as drying occurs.

**Figure 8.**
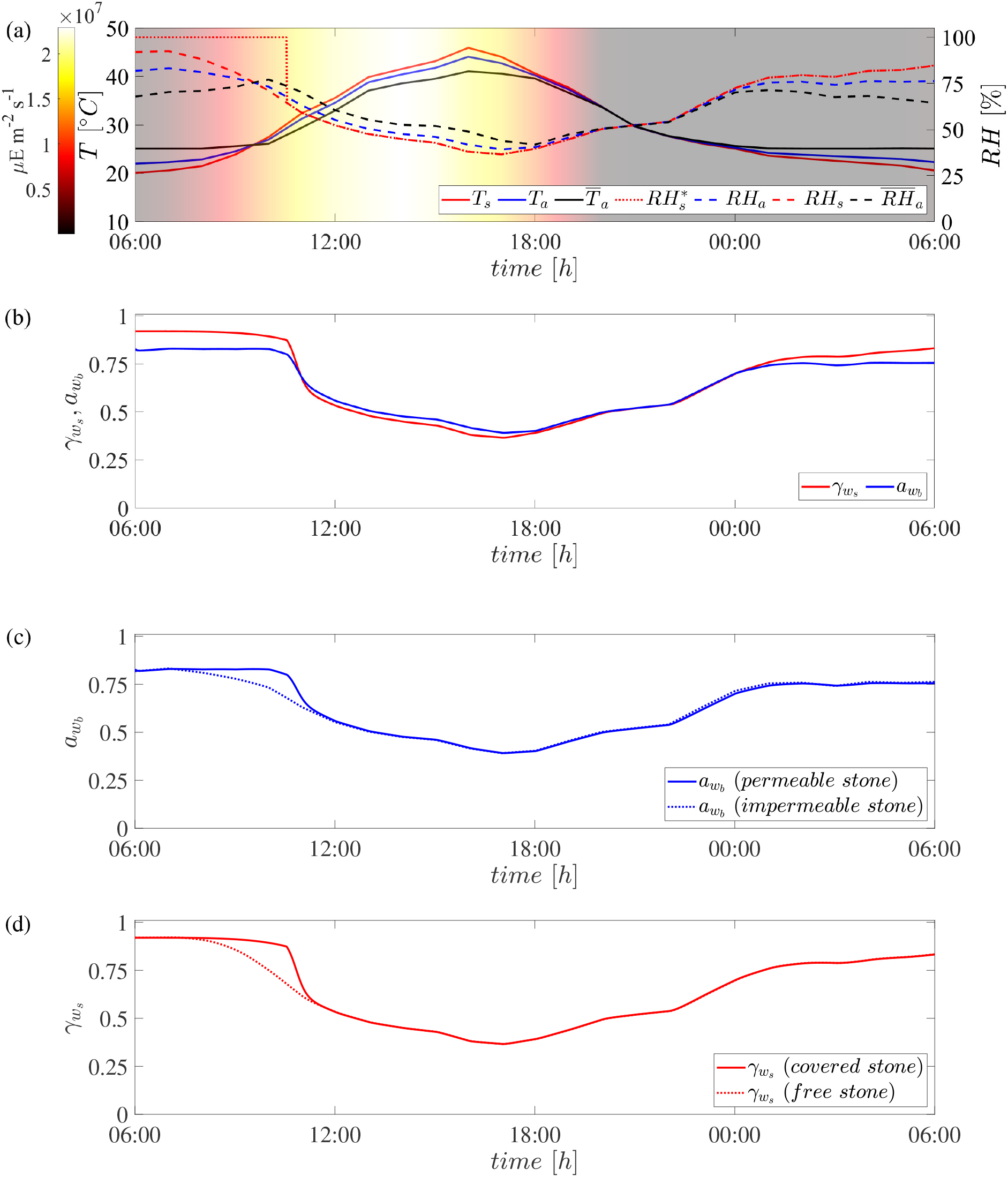
(a) One Day profile of temperatures and relative humidities in the compartments of the SAB-stone system, and light intensity, on a summer day (June 18 *−* 19, 2016) at the Thomas Jefferson Memorial, Washington D.C., USA [41]. (b) Scenario 1 stone pore water activity coefficient 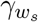 and SAB water activity 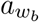 in the SAB-covered stone. Note water activity is suitable for biolofical activity only during part of the day. (c) Scenario 2 - comparison of SAB water activities *a*_*w*_*b* for the SAB-covered stone for the permeable and impermeable stone cases. Added effect of water storage in stone pores increases SAB water activity during early daylight hours. (d) Scenario 3 - comparison of stone pore water activity coefficient 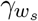 SAB and SAB-free cases. Note that the main differences in (b)-(d) are in the morning hours, when water conditions are most optimal, and which are the most critical for SAB activity.

We consider three specific model scenarios for comparison:

– **Scenario 1**: A reference configuration in which the system consists of both a subaerial biofilm and the underlying permeable stone.
– **Scenario 2**: A system composed of a subaerial biofilm but the underlying stone is supposed impermeable (implemented, practically, by setting *σ*_*ex*_ = 0), in which SAB water activity is evaluated under the assumption of instantaneous evaporation/condensation (high *σ*_*b*_).
– **Scenario 3**: A system consisting of a bare (permeable) stone, i.e., no SAB, in which water dynamics within the pore network are driven solely by interactions with the external ambient air (implemented, practically, by considering only the equations for the stone layer and setting *η* = 1 and *σ*_*ex*_ = 0).

In all simulations, initial conditions for (12) are set such that the system is in equilibrium with the environmental conditions at *t* = 0. Model parameters used for the computations are collected in Table 1.

**Table 1.** Model parameters used in the numerical simulations.

| Parameter | Definition | Unit | Value |
| --- | --- | --- | --- |
| $\phi_s$ | Stone porosity | % | 8.91 <sup>(a)</sup> |
| $V_s$ | Total volume of the limestone layer | $mm^3$ | $75 \times 23 \times 10$ <sup>(a)</sup> |
| $I_s$ | Interface between air and water in stone pores | $mm^2$ | 1.61 <sup>(a)</sup> |
| $I_b$ | Interface between ambient air and SAB | $mm^2$ | $75 \times 23$ <sup>(a)</sup> |
| $I_{ex}$ | Interface between SAB and stone layers | $mm^2$ | $75 \times 23$ <sup>(a)</sup> |
| $\sigma_s$ | Evaporation and condensation coefficient at $I_s$ | — | $2.7 \cdot 10^{-4}$ <sup>(a)</sup> |
| $\sigma_b$ | Evaporation and condensation coefficient at $I_b$ | — | $5 \cdot 10^{-6}$ <sup>(a),(b)</sup> |
| $\sigma_{ex}$ | Evaporation and condensation coefficient at $I_{ex}$ | — | $5 \cdot 10^{-6}$ <sup>(a),(b)</sup> |
| $\tilde{a}_{wb}$ | Threshold value of water activity | — | 0.8 <sup>(a)</sup> |
| $\tilde{S}_{ws}$ | Threshold value of water saturation in the stone | % | 30 <sup>(a)</sup> |
| $\tilde{S}_{wb}$ | Threshold value of water saturation in the SAB | % | 60 <sup>(a)</sup> |
| $\delta$ | Coefficient of the equation for $\gamma_{ws}$ and $a_{wb}$ | — | 20 <sup>(a)</sup> |
| $\varepsilon$ | Coefficient of the equation for $\eta$ | — | 15 <sup>(a)</sup> |
| $l_b^*$ | Average thickness of SAB | $\mu m$ | 40 <sup>(a)</sup> |
| $l_{bl}$ | Thickness of the thermal boundary layer | $cm$ | 1 <sup>(c)</sup> |
| $k_a$ | Thermal diffusivity of air | $m^2 s^{-1}$ | $2.2 \cdot 10^{-5}$ <sup>(c)</sup> |
| $k_b$ | Thermal diffusivity of SAB | $m^2 s^{-1}$ | $1.43 \cdot 10^{-7}$ <sup>(c)</sup> |
| $a$ | Magnus formula coefficient | $Pa$ | 610.94 <sup>(d)</sup> |
| $b$ | Magnus formula coefficient | — | 17.625 <sup>(d)</sup> |
| $c$ | Magnus formula coefficient | $^{\circ}C$ | 243.04 <sup>(d)</sup> |

(a) discussed in Section 2 and 3.
(b) varied in the numerical simulations.
(c) from [40].
(d) from [2].

Figure 8b displays time variations of stone pore water activity coefficient 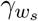 and SAB water activity 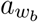 for Scenario 1, permeable stone with SAB (reference case). The daily temperature variation causes a pronounced variation in relative humidity. Consequently, the water content in both the stone and the SAB undergoes slight variations over the 24-hour period (not shown), which are nevertheless sufficient to produce more noticeable oscillations in the associated SAB water activity and in the stone pore water activity coefficient. Note the curves separate significantly at night; during periods when the water activity in the SAB exceeds 0.8, the stone matrix behaves as a closed system so that relative humidity within the stone pores 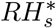 reaches high values - often approaching saturation. The principal observation, though, is that water activity remains flat and relatively high until late morning.

Figure 8c compares results of Scenario 2 (SAB with impermeable stone) with the reference case (SAB with permeable stone). SAB water activities closely track each other and the ambient air relative humidity, except for a period during the late morning. When accounting for water exchanges between the SAB, the stone, and the ambient air, we see water activity exceeds the values observed in the impermeable stone case during late morning, remaining above 0.8 for a longer period. Indeed, when stone behaves as a closed system, evaporation from the pores is delayed. As a result, water in the stone pores acts as a reservoir and moves to the biofilm to compensate for atmospheric evaporation. This effect, in the model, roughly doubles the time when both water activity and light conditions are simultaneously favorable for growth.

Figure 8d compares the stone pore water activity coefficient of Scenario 3 (permeable stone with no SAB) with that of the reference case (permeable stone with SAB). We note that SAB presence increases stone water content, but only modestly, suggesting that SABs do not impact stone water conditions significantly. It is important to note, though, that our model is not designed to consider other possible impacts, ignoring factors like salinity, temperature, and heterogeneity.

## 5 Discussion

Microbial colonization of air-exposed solid substrates is dependent on the presence of free, liquid water, which is essential for metabolic activity. In dry environments, SABs are sustained by their ability to obtain, and retain, liquid water. Characterization of that process is central to understanding these communities. However, in the field and the lab, SAB-stone water dynamics are difficult to measure to the point that modeling may be essential. This study presents a basic model of the primary physical and chemical processes governing water dynamics in systems composed of interacting subaerial biofilms and porous stone/masonry substrates. We argue that water activity within SABs can be impacted in a significant way by its underlying permeable substrate. In particular, a framework is proposed to estimate water availability in SABs by accounting for evaporation and condensation within the SAB-stone system, and for the water exchange between the biofilm and the substrate. These mechanisms are fundamentally thermodynamic and so are expressed through gradients of water chemical potential in combination with the water affinities of both the SAB matrix and the stone pore network. Together with a notion of closed and open SABs, these considerations are already able to successfully explain laboratory measurements of SAB-limestone interactions.

Model results suggest that the presence of SABs affects the water evaporation process in a subtle but significant manner by delaying evaporation and degradation of water activity during the critical morning hours when conditions for metabolic activity are optimal. This aligns with observations and supports the idea that, in the morning, SABs act as sponges, drawing water from the stone pores to compensate for atmospheric evaporation [1, 44, 46]. That is, basic thermodynamics supports the idea that the solid substrate actively influences and regulates water availability, serving as both a moisture source and buffer and supplying water to the SAB when needed [21, 43, 46, 49]. In this way, SABs can benefit from the interaction with a permeable rock substrate, as SABs themselves can be too thin to maintain sufficient water on their own.

We note here that while results are intended to apply generally to stone and masonry materials, we have only parameterized and validated using SAB on limestone data. We expect that the generality of the model will allow parameters associated with limestone to be replaced with those of other materials, allowing the model to be adopted to investigate how different solid substrates influence water transport and activity within SABs. Nevertheless, the function representing the activity of water in the porous medium under consideration could be re-visited, as it depends on the material properties. The model itself is a rather rough one, in line with the limited availability of observational data which does not allow a more detailed one, and we also expect it can be revised and improved with new observations.

## A Thermodynamic description of chemical potential in a liquid-vapor system

Evaporation and condensation are generally nonequilibrium processes, characterized by the net transfer of molecules across a liquid-gas interface into the gas phase or liquid phase, respectively. A fundamental concept in describing such processes is the chemical potential *µ*_*j*_ of a species *j*, the free energy available per mole of that species to perform work [22]. Chemical potential depends on multiple factors, including species concentration and partial pressure, electrical potential, and gravitational effects [22, 29] and can be viewed as an intrinsic property of species *j*, indicating how it will respond to changes in the system. For example, if species *j* undergoes a transition from state *A* to state *B*, its chemical potential changes from 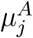 to 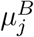. In particular, if 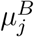 is less than 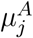, then the free energy per mole of species *j* decreases.

The condition for a spontaneous chemical change is an opportunity to decrease the chemical potential. This principle has direct implications when analyzing fluxes between different regions. A difference in chemical potential, 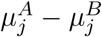, serves as a *driving term* for the movement of species *j*: the greater this difference, the faster the transition will occur. By comparing chemical potentials on opposite sides of a boundary or interface, we can assess whether a species is in equilibrium or predict the direction and the velocity of the movement. If *µ*_*j*_ is equal on both sides, no net movement is expected. Conversely, any difference implies a tendency for species *j* to move toward the region with lower chemical potential.

### Liquid water chemical potential

Here we assume that the chemical potential of liquid water *µ*_*w*_ depends mainly on its chemical composition and interactions, and is given to good approximation by

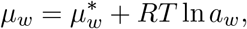

where 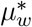 is the chemical potential of liquid water in a standard (ideal) reference state, and *a*_*w*_ is the water activity, which quantifies the thermodynamic availability of water and accounts for the interactions between solvent and surrounding particles. In a solution, molecules of water interact with solvent, and undissolved, molecules. These interactions influence water behavior, and are captured through the water activity, which is defined as *a*_*w*_ = *γ*_*w*_*N*_*w*_, where *N*_*w*_ is water mole fraction and *γ*_*w*_ is the activity coefficient. In an ideal solution *γ*_*w*_ = 1, meaning that the interactions between water and surrounding molecules are equivalent to those with pure ideal water. In non-ideal solutions, 0 *< γ*_*w*_ *<* 1 generally.

### Water vapor chemical potential

Water molecules in an aqueous solution are constantly exchanged with the surrounding gas phase (when present), with evaporation and condensation occurring simultaneously. The rates of these two processes depend on the chemical activities of water in both the liquid and gas phases. At equilibrium, the rates of evaporation and condensation are equal. Under this condition, the adjacent gas phase holds the maximum possible amount of water vapor for a given temperature, i.e., the gas phase is saturated. The partial pressure of water vapor in equilibrium with liquid water is known as the water vapor saturation pressure 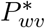. The chemical potential of water in the vapor phase *µ*_*wv*_, which depends to good approximation in many circumstances only on the water vapor partial pressure *P*_*wv*_ [22, 29], can be approximated as

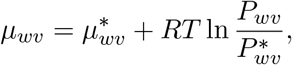

where 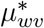 is the chemical potential of water vapor in a standard (ideal) reference state. At relative humidity of 100%, air in contact with pure water contains the maximum vapor pressure possible at that temperature. Note that saturation pressure increases significantly with temperature.

In practice, when evaluating mass transfer between two regions, we are interested in differences in chemical potential rather than their individual values. At local equilibrium, with liquid and gas phases at the same temperature, the reference chemical potentials for the liquid and vapor phases are equal, i.e., 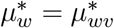 and thus

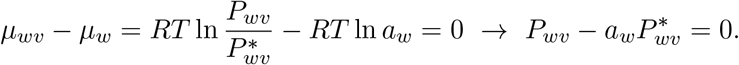

We can observe three possible situations:

– Negative *driving stress* 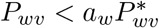: this means that the water vapor partial pressure is lower than the water vapor saturation pressure, and the relative humidity is less than the water activity. This indicates that the system is not yet saturated and it is in a state where the liquid phase can evaporate into the vapor phase (*µ*_*wv*_ *< µ*_*w*_).
– Positive *driving stress* 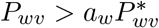: this means that the water vapor partial pressure is higher than the water vapor saturation pressure, and the relative humidity is greater than the water activity, suggesting that net condensation of water vapor into the liquid phase will occur in order to reach equilibrium (*µ*_*wv*_ *> µ*_*w*_).
– Vanishing *driving stress* 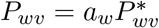: this means that there is a dynamic equilibrium between the liquid and vapor phases, where the rate of evaporation equals the rate of condensation, and the system is in a stable state. In other words, no net phase change occurs when the water vapor partial pressure equals the product of the water vapor saturation pressure and the water activity (*µ*_*wv*_ = *µ*_*w*_).

## B Environmental parameters evaluation

**Water vapor saturation pressure** 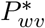 depends on the temperature *T* and can be approximated using the empirical Magnus formula

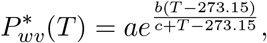

with coefficients *a* = 610.94 Pa, *b* = 17.625, *c* = 243.04 ^*°*^C [2].

### Temperature and relative humidity on the SAB surface

To account for the presence of a thermal boundary layer, it is necessary to evaluate the temperature and relative humidity at the biofilm-boundary layer interface. These quantities can be computed using the relationships proposed in [41]

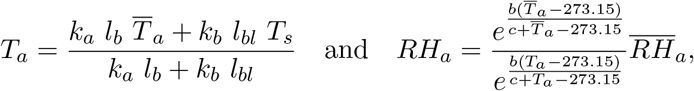

where *k*_*a*_ and *k*_*b*_ are respectively the thermal diffusivities of air and the SAB, *l*_*b*_ is the SAB thickness, *l*_*bl*_ is the thickness of the thermal boundary layer, 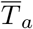 and 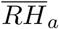 are the temperature and relative humidity of external ambient air, and *T*_*s*_ is the stone temperature.

### Estimation of relative humidity in the ambient air

Weather data from the marble roofs of the portico at the Thomas Jefferson Memorial used in this work include stone temperature and relative humidity, ambient air temperature, and light intensity. For this reason, ambient relative humidity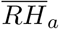 is needed. It is estimated following the approach described in [41]

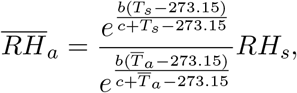

where the temperature and humidity data from the stone surface together with the ambient air temperature are used.

## Acknowledgments

The authors would like to thank Federica Villa for helpful discussions.

The authors would also like to thank Bettina Buttaro, Jean-David Grattepanche, Judith M. Jacob, and Alberto Tenore for their assistance.

The authors would like to thank the US National Park Service. The authors acknowledge support from NSF/DMS 1951532.

This paper has been performed under the auspices of the G.N.F.M. of I.N.d.A.M.

